# Semantic relation priming is not constituent-specific: Evidence from ERP

**DOI:** 10.1101/2021.11.10.468040

**Authors:** Xiaofei Jia, Changle Zhou

**Author notes:** Send correspondence to Changle Zhou, Department of Cognitive Science, Xiamen University, Xiamen, China,.

## Abstract

It is humankind’s unique wisdom to compose a limited number of words together through specific rules to convey endless information. Researchers have found that this composition process also plays a vital role in the comprehension of compounds. The specific manifestation is relation priming; that is, the previously used relation will promote subsequent word processing using the same relation. This priming phenomenon is bound to morpheme repetition (modifier or head). This study combines a self-paced priming paradigm with electrophysiological technology to explore whether relation priming will occur without sharing morphemes and its time course. We found that relation priming can occur independently of morpheme-repetition, which shows an independent representation of relation information. And it has been activated at a very early stage (about 200ms). As the word processing progresses, this activation gradually strengthens, indicating that the relation’s role is slowly increasing in the process of compound word recognition. It may first be used as a kind of context information to help determine the constituent morphemes’ meaning. After the meaning access of the constituent morphemes, they begin to play a role in the semantic composition process. This study uses electrophysiological technology to precisely describe the representation of relation and its time course for the first time. Which gives us a deeper understanding of the relation priming process, and at the same time, shed light on the meaning construction process of compounds.

## 1. Introduction

In recent decades, a focus of controversy about compounds is whether it is represented as the whole word or decomposed into constituents in a mental dictionary. The traditional view is that compounds are processed as a whole. When we acquire a compound, its representation is formed in the mental lexicon. When we reencounter it in reading, we can quickly extract its meaning (Butterworth, 1983; Bybee, 1995). This view sounds very reasonable, and it is precisely because of this rapid extraction process that each of us can read quickly and effortlessly. However, if each word has its mental representation, we have to have an astronomically sizeable mental vocabulary, which will be a huge memory burden. Therefore, in recent years, there appeared much evidence to support the view of decomposition. It is believed that compounds are broke down into its components and stored in the mental dictionary. When comprehending a compound word, we need to figure out its constituent morpheme’s meaning first and then put them together to construct the meaning of the whole word (Libben & Jarema, 2004; McKinnon, Allen, & Osterhout, 2003; Taft & Forster, 1975). In this way, only a limited number of morphemes needs to be stored in the mental dictionary, and innumerable concepts can be constructed. This seems to be a more economical way.

In recent years, researchers have conducted many experiments around this composition process and found that not only the constituent morphemes but even the relation connecting these morphemes play a vital role in the composition process (Gagné & Spalding, 2009). This seems to be easy to understand. Other creatures can only use a limited single vocabulary to communicate information and express ideas. While the world is all-encompassing and limited available vocabulary is difficult to cover everything. Only humans have developed the ability to compose different terminology according to a specific structure, just like building a castle with various building blocks. Two-morpheme compounds can be regarded as the simplest language combination, and relation can be viewed as the simplest combination rule. Studying the compounding process of two-morpheme compounds helps to clarify the basis of the more complex combination ability.

The relation can be divided into two categories, grammatical relation and semantic relation. In English compounds, the modifier-noun structure is the dominant structure, while in Chinese, there are four other grammatical structures. However, structure priming has been found in English and Chinese (Gagné & Shoben, 1997; Ji & Gagné, 2007; Jia, Wang, Zhang, & Zhang, 2013). The relation we mentioned in this article refers to the semantic relation. For example, Gagne et al. found that the target word of modifier + noun structure can be facilitated by the prime word with the same semantic relation. For example, the participant would feel *student vote* (vote by a student) is easier to understand than *student car* (car of a student) if they have just watched *student accusation* (accusation by a student) (Gagné, 2000, 2001; Gagné & Shoben, 2002). Similar phenomena have also been found in Chinese.

However, there is no consensus on the representation of relations. Gagne et al. found that the relation’s effects are constituent-specific (modifier) and did not observe general relation priming. Only when the priming word and the target word share morphemes will the relation priming effect occur (Gagné, Spalding, & Ji, 2005). But Estes came to the opposite conclusion. He believes that relation priming is essentially a unique mode of activation; regardless of whether there is morpheme repetition, the facilitated effect of relation will be manifested. The reason why Gagne’s experiments failed to find the independent representations of relation maybe because the semantic classification standards they adopted were rather vague, which did not capture the essence of the relation. Moreover, they could not distinguish between the same relation and the different relations well (Estes & Jones, 2006). In Chinese, researchers also found such relation priming in their experiments (Ji & Gagné, 2007; Jia, Wang, Zhang, & Zhang, 2013). Moreover, they found the mode of relation priming is affected by language characteristics. Regardless of the repeated morpheme is a modifier or head, it can trigger relation priming. However, their experiment only sets morpheme repetition conditions, so it is impossible to answer whether relation priming can occur independently of morpheme repetition.

Raffray et al. found that relation priming can occur without morpheme repetition, but morpheme repetition can boost relation priming (Raffray, Pickering, & Branigan, 2007). Therefore, we speculate that relation priming can occur without morpheme repetition. Since the semantic classification used by Gagne et al. is not typical enough, only relatively strong relation priming is found; and the relation priming is much weaker when there is no morpheme repetition, so they failed to detect it. Chinese is a meaning-spelling language. One person only needs to master seven or eight thousand characters(i.e., semantic bases). The vocabulary formed by combing these bases can meet daily needs(Zhang, 2011). This feature determines that compounding is its primary word-formation method.

Moreover, Chinese only has morpheme boundaries but no word boundaries, so that it can be combined more freely. So the Chinese should have greater freedom of combination and higher productivity to meet their language characteristics. Independent representation of relational seems to be a need. For the above reasons, we use Chinese materials, and these materials only contain typical instances of each relation (the consistency of the rate between subjects reached above 95%). If the above speculation is correct, then the relation priming should also appear in the absence of morpheme repetition. The facilitate effect under the repeated condition should be more significant than that of the non-repeated condition.

We set up morpheme repetition conditions (modifier or head, corresponding to MS, HS) and morpheme non-repetition conditions(NU) in the following experiments. We expect that relation priming will appear under both conditions. That is, under the conditions of MS, HS, and NU, the same relation will have better behavior performance than the different relations. The relation priming under the morpheme repetition condition is greater than that of the non-repetition condition. That is, the difference between the same relation and the different relation is more significant under the MS and HS conditions than under the NU condition.

Jia’s research found that the time when relations begin to take effect is later than the time when constituent morphemes are semantically activated (Jia et al., 2013). Because relations’ starting to work is on behalf of the beginning of the meaning composition process, so they always occur after the semantic activation of constituent morphemes. But this is limited to the relation associated with the modifier because the relation priming they found occurs under the modifier-repeated condition (Jia et al., 2013). Which is currently the only ERP study on the time course of relation priming. We want to throw light on the relation priming’s time course under head-repeated and no repetition condition(if there is a relation priming). Therefore, in addition to modifier-repetition conditions, the current research also sets sharing head conditions and non-sharing morpheme conditions.

Some researchers believe that modifier is the first morpheme, which is more critical for whole-word processing than the last morpheme (Zhang et al., 2012). Some researchers believe that for the modifier + head structure, the head represents the conceptual category to which the entire word belongs, while the modifier only modifies details, so the head is more critical for whole-word processing (Gagné & Shoben, 1997; Manouilidou, Ralli, & Kordouli, 2012). In the head-repeated condition, the rough meaning of the whole word would be activated when the head was primed. In this case, would the relation that only serves as a modification still work? If it does, would the pattern be the same as that in the modifier-repeated condition? If it was different, what pattern would it show? And if the relation priming exists when there are no repeated morphemes, what pattern would it lead? Should it be activated at the same time as or after the constituents? If the activation occurs simultaneously with constituents, should it start at the lexical or semantic level? Therefore, besides the N400 time window (relation priming associated with modifier occurs in this window), we would also examine whether the relation activated in an earlier period, such as the morphological processing stage.

The central-parietal N200 is an ERP component induced by Chinese two-morpheme compounds recently discovered by Zhang, which appears 200 ms after the stimulus onset, and centered on the central-parietal area (Zhang et al., 2012). At present, it is considered a component related to the processing of Chinese orthography. If the relation is activated at this stage, the same relation and different relation conditions should show differences in the N200. On the contrary, there will be no difference.

## 2. Method

### 2.1 Participants

Thirty-two healthy college students participated in this experiment voluntarily. All of them were native Chinese speakers, right-handed, aged between 18-25 years (average 21.4 years), with normal or corrected to normal vision. All participants read and signed the written informed consent following a research protocol approved by the IRB board of Xiamen University before the experiment.

### 2.2 Research methods

The target includes 72 noun-noun compounds with a “modifier + head” structure(Some of them were selected from Ji’s research (Ji & Gagné, 2007)). Each target (e.g., *手表* meaning wristwatch) has six primes corresponding to the six conditions: 1. Sharing the modifier and the relation with the target (MSRS, e.g., *手链* meaning hand chain); 2. Sharing the modifier with but differing in the relation from the target (MSRD, e.g., *手背* meaning hand back); 3. Sharing the head and the relation with the target (HSRS, e.g.,*怀表* meaning pocket watch); 4. Sharing the head with but differing in the relation from the target (HSRD, e.g., *秒表* meaning stopwatch); 5. Sharing no constituent but have the same relation with the target (NURS, e.g., *颈环* meaning neck ring); 6. Sharing neither constituent nor the relation with the target (NURD, e.g., *丝袜* meaning silk stockings).

According to the frequency dictionary of Cai et al. (Q. Cai & Brysbaert, 2010), the mean frequency of the target is 0.94 occurrences per million. There was no statistical difference in mean frequency across six conditions (MSRS: 0.91, MSRD: 0.79, HSRS: 1.01, HSRD: 0.89, NURS: 0.85, NURD: 1.03). Using a 5-point scale (1 means completely irrelevant, 5 means highly correlated), 15 people from the same subject population but not participate in this experiment rated the prime and target’s semantic relatedness. There was no statistical difference between the RS and RD conditions at the three morpheme levels (2.42 vs.2.65, p>0.5; 3. 02 vs.2.96, p>0.5; 2.04 vs.1.97, p>0.5). Using a 5-point scale (1 means not in the same category, 5 means in the same category), another 15 people rated the category relatedness between the prime and target, that is, judging the likelihood of the prime and target coming from the same semantic category. And there was no statistical difference between the RS and RD conditions at the three morpheme levels (2.13 vs.2.24, p>0.5; 2.84 vs.3.05, p>0.5; 1.78 vs.1.96, p>0.5).

One list of 432 filler pairs was also constructed. In half of the word pairs, the prime was interpretable, but the target was nonsense. In the other half, both the prime and the target were nonsense. Matched with the test pairs, in 2/6 (144 pairs) of the total filler pairs, the first morpheme was repeated. In another 2/6 (144 pairs), the second morpheme was repeated, and the remaining 2/6 do not have repeated morphemes. All test pairs and filler pairs are divided into six parts, forming six versions. Each version contains 72 test pairs and 72 filler pairs. Each target appears only once in each version, and its corresponding six primes appear in the six versions, respectively. Each participant used one version for a test.

We adopt a sense-nonsense judgment task and a priming paradigm. Participants sat in a sound-attenuated and dimly lit room facing the screen with a distance of 0.7 m while placing the index fingers of the left and right hands on the F and J keys of the keyboard, respectively. They were asked to look at the fixation in the center of the screen and relax. At the beginning of each trial, “Ready?” appeared in the center of the screen. When they were ready, they can press the Q key to initiate each trial. First, the prime appeared, and the participant judged whether the word is meaningful or not and respond by pressing the corresponding key (F or J) as quickly as possible. After they responded, the message “Ready?” appeared again, and participants press the Q key to display the target. Likewise, participants indicate whether the target is meaningful or not by pressing the F or J key. Two adjacent trials are a unit, where the first presents the prime, and the second shows the target, but this pair structure was not made known to them. The self-paced presentation gave participants enough time to process each word and ensure that they could fully understand each word and responded according to its semantic sensibility.

### 2.3 EEG recording and data analysis

Participants were required to avoid unnecessary head movement and control blinking during the stimulus presentation and responding during recording sessions. Using NEUROSCAN 4.5 with 64 -channel Ag/AgCl electrode caps, the electrodes are placed according to the extended 10-20 system, the band-pass filtering range is 0.1-70 Hz, and the sampling rate is 500 Hz. The impedance in all electrodes was less than 7 kΩ. The electrodes for recording the vertical EOG were located above and below the left eye, and the electrodes for recording the horizontal EOG were located on the left and right lateral orbital rim. Taking the tip of the nose as a physical reference, the original EEG was recorded continuously and was re-referenced offline to the mean of the bilateral mastoids. EEGLAB 14.1.1 was used for data analysis. Epochs were computed from 200ms before to 800ms after the stimulus onset, and the baseline correction was made between −200 to 0ms. The offline band-pass filtering was applied between 0.1-30 Hz, and independent component analysis (ICA) was used to remove Ocular artifacts. Incorrect responses and responses with amplitude greater than ±75 μv are excluded from the superimposed average. The data rejected due to artifacts is 8.5%.

## 3. Results

### 3.1 Behavioral results

For the response time data, we used LmerTest package of R software to fit a mixed-effects model with logarithmic response time as the dependent variable, morphemes (MS, HS, NU) and relations (RS, RD) as fixed factors, and subject and item as random factors (Baayen, Davidson, & Bates, 2008; Kuznetsova, Brockhoff, & Christensen, 2017; Team, 2013). We found that morphemes have a main effect, F (2,2287)=6.99, p<0.001, and the relation also has a main effect, F (1,2276) =7.19, p<0.01, but there is no interaction between them, F (2,2276) = 0.97, p>0.1. Contrast analysis showed that the response time of NU conditions was significantly slower than that of MS and HS, while there was no significant difference between MS and HS (MS vs. NU: 794 vs. 850 ms, t (2290) = −2.60, p<0.01; HS vs. NU: 771 vs. 850ms, t (2288) = −3.65, p<0.0005; MS vs. HS: 794 vs. 771 ms, t (2289) = −1.05, p>0.1). The response time of the RS condition is significantly faster than that of the RD condition (RS vs. RD: 850 vs. 897ms, t (2278) = −2.68, p<0.01). The logistic regression method was used to analyze the accuracy data. Because the accuracy reached the ceiling (above 98%), all the analyses were statistically non-significant.

### 3.2 ERP results

Figure 1 shows the average ERP response on the representative electrodes using different colors for the six conditions(MSRS, MSRD, HSRS, HSRD, NURS, and NURD). Waveforms at electrode Cz are highlighted in Figure 2 for clarity. It can be seen that the above six conditions all elicited a clear N400. We take a 300-450ms time window and do a morpheme (3) *relation (2) repeated-measures ANOVAs of the averaged amplitude over 9 electrodes (FC1, FCz, FC2; C1, Cz, C2; CP1, CPz, CP2). We found that morphemes have a main effect, F (2,155) =11.54, p<0.0001, and the relation also have a main effect, F (1,155) =4.01, p<0.05, but there is no interaction between them, F (2,155) =0.03, p>0.5. Contrast analysis showed that, there is a significant reduction in amplitude of MS and HS relative to NU, and there was no significant difference between them (MS vs. NU: 0.52 vs. −1.07 μV, t (157) = 4.43, p<0.0001; HS vs. NU: 0.32 vs. −1.07 μV, t (157) = 3.89, p<0.0005; MS vs. HS: 0.52 vs. 0.32 μV, t (157) = −0.55, p>0.5). RS also showed a significant reduction in amplitude compared with RD conditions (RS vs. RD: −1.07 vs. −1.66 μV, t (157) = −2.02, p<0.05). Because the relation information is our focus, to show it more clearly, we show the relation effects of MS, HS, and NU conditions on behavior, N200, and N400 separately, as shown in Figure 3.

Figure 1. should be inserted here

**Figure 1.**
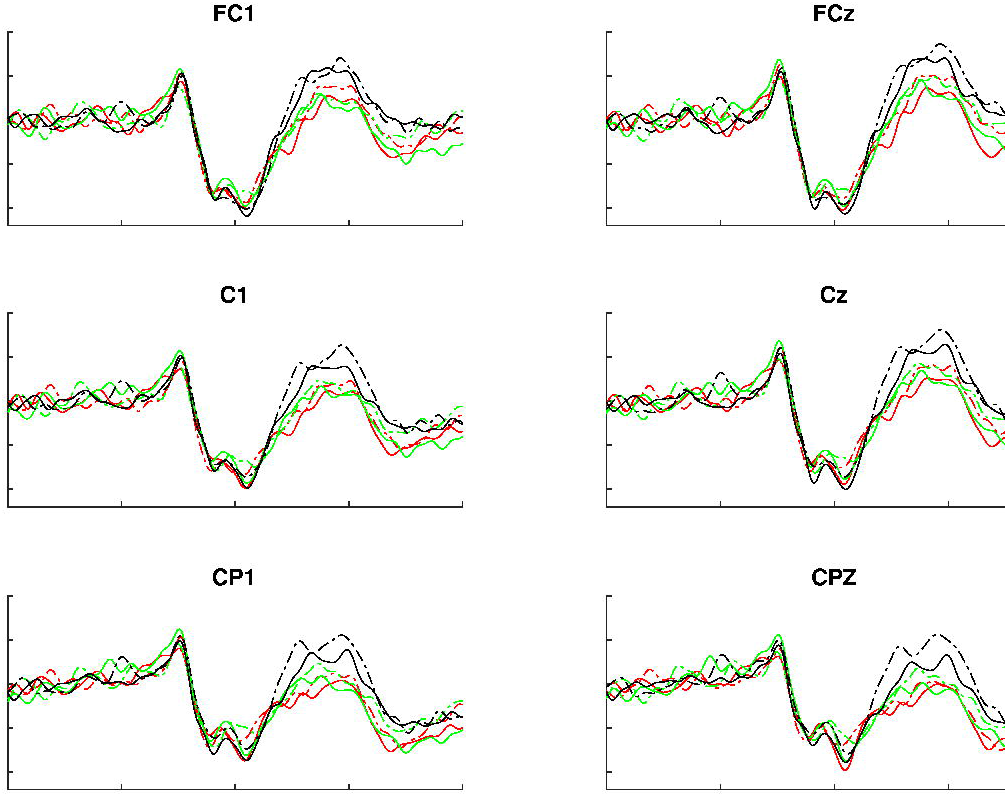
Grand average ERP waveforms for all conditions in 9 representative electrodes. MSRS: prime Sharing the modifier and the relation with the target; MSRD: prime Sharing the modifier with but differing in the relation from the target; HSRS: prime Sharing the head and the relation with the target; HSRD: prime Sharing the head with but differing in the relation from the target; NURS: prime Sharing no constituent but have the same relation with the target; NURD: prime sharing neither constituent nor the relation with the target.

Figure 2. should be inserted here.

**Figure 2.**
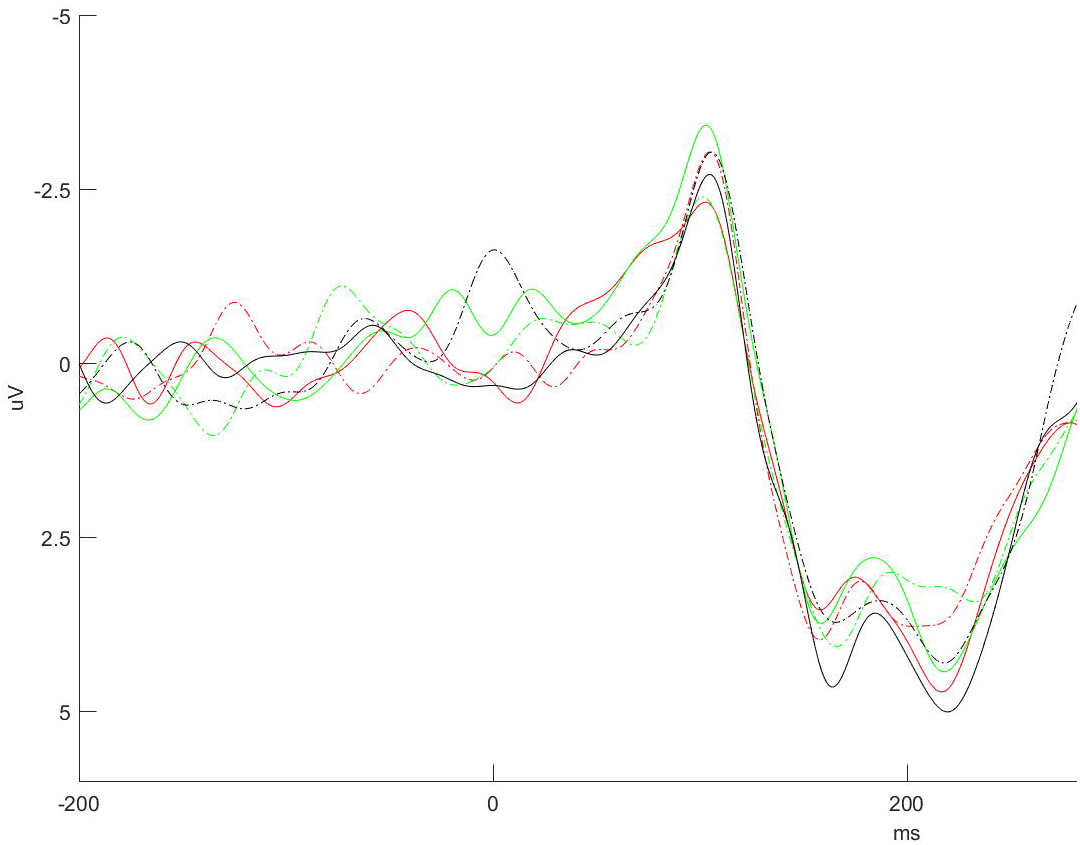
Electrode Cz from Figure 1 highlighted for clarity. Legends are the same as in Figure 1.

Figure 3. should be inserted here

**Figure 3.**
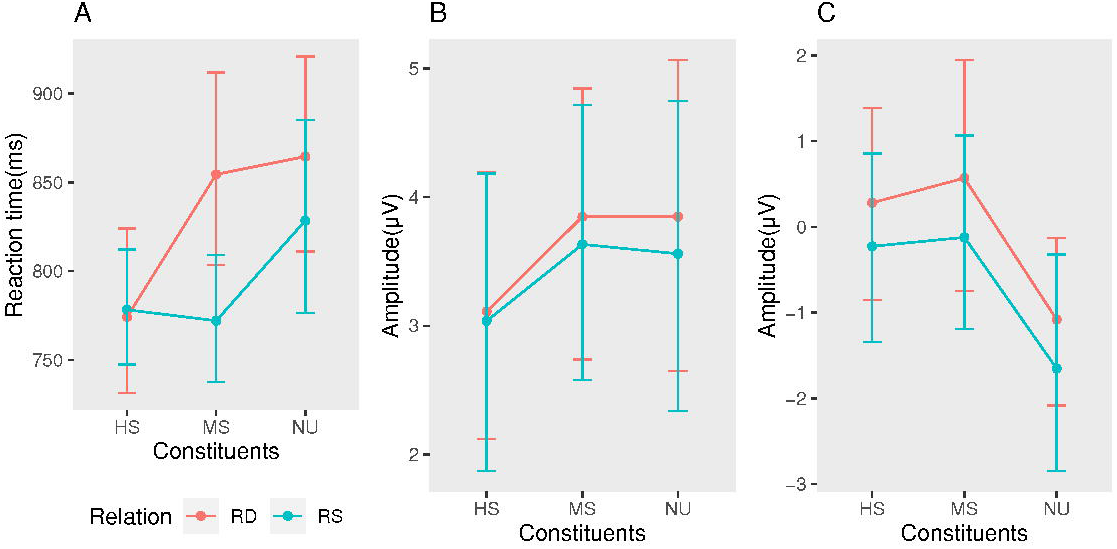
Relation effects on Reaction time(3-A), N200(3-B), and N400(3-C). RS: prime sharing the same relation with the target; RD: prime have different relation from the target.

To test whether this relation effect appeared in the earlier period(N200), we took a time window of 185-207ms and performed a morpheme (3) * relation (2) repeated-measures ANOVAs on the average amplitude of the nine electrodes mentioned above. No main effects were found in morphemes and relations (F (2,155) =1.93, p>0.1; F (1,155) =0.38, p>0.5), and there was also no interaction between them (F (2,155) =0.04, p >0.5). However, as shown in Figure 3-B, although not reached significance, there is a trend of relation effects. That is, RS conditions have increased amplitude relative to RD in MS and NU, except for HS.

## 4. Discussion

As the behavioral results indicated, MS and HS show better behavioral performance than NU because there is a repetition of the first morpheme between the prime and target, reflecting the repetitive priming effect of the morpheme and the semantic priming at the whole word level. Under all these three conditions, RS has better behavior performance than RD because there is a repetition of the relation between the prime and target, reflecting the effect of relation priming. It’s just that the relation priming effect under the constituent-repeated condition (MS and HS) is not greater than that of the constituents’ non-repeated condition (NU), which does not meet our expectations. We will explain the reason later.

In the EEG results, the results of N400 is consistent with the pattern of behavioral consequences. MS and HS have a significant reduction in amplitude relative to NU, reflecting the repetitive priming of morphemes and semantic priming at the whole word level, which is consistent with the classic N400 effect. What is important is that at these three levels of NU, MS, and HS, RS has a significant reduction in amplitude relative to RD, showing an apparent relation priming effect. The relation priming at the morpheme-repeated level (MS and HS) repeats Ji and Jia et al. (Ji & Gagné, 2007; Jia et al., 2013). The new result we found is that the relation priming can also occur independently (NU) without sharing morphemes. It shows that relation can be independently represented and does not have to be bound to morphemes. This is in line with the prediction of the Transformational model of relation representation (Leech, Mareschal, & Cooper, 2008). They believe that the relation is represented as a unique activation mode triggered by multiple inputs. Initial exposure to a situation will trigger a relation. Then this relation can be applied to a new situation for analogy. In this experiment, regardless of whether there is a repetition of constituents, the activation pattern triggered by prime can be mapped to target, thereby facilitating its processing.

Compound words are equivalent to the simplest language composition. The composition rule of sentences is grammar, and the composition rule of compound words is the relation. The relation could be regarded as the simplest composition rule. Therefore, relation priming could be considered as a kind of structural priming. Martin Pickering et al. found a syntactic priming effect in a language production experiment. The speaker tends to re-use a specific syntactic structure previously used. For example, suppose the speaker heard a subject-predicate sentence before. In that case, he is more inclined to use subject-predicate sentences rather than prepositional-objective sentences in his following speech, although prepositional-objective sentences can also express the same meaning. When the critical verbs used in the sentence that has been experienced and the sentence that needs to be expressed are the same or have a similar meaning, the tendency to repeat the syntactic structure is particularly strong (Pickering & Branigan, 1998). (This is very similar to relation priming. Only when the prime and target use the same relation and the same morpheme, the relation priming is particularly strong. Later, another experiment found that the syntactic priming exists independently of the repetition of the concepts (such as verbs) expressed in the sentence (ZG Cai, Pickering, & Branigan, 2012; Huang, Pickering, Yang, Wang, & Branigan, 2016; Yu Zhou & Zhang Qingfang, 2020). How similar is this to the discovery process of relation priming! In the beginning, we thought that relation priming would only occur in the case of morpheme repetition, but now we have discovered that it could exist independently of morpheme repetition! We believe that the essence of syntactic priming and relational priming is the same; both are the embodiment of structure priming at different levels. One is at the sentence level, and the other is at the word level. And they are essentially manifestations of the role of composition rules (structural mapping) in language processing. The independent representation of the composition rule (i.e., structure, relation) adds new evidence for compounds’ decomposition view.

Above, we have proved that relations are represented independently. Next, we will look at the time course of relation priming.

Combine the ERP results to see the time course of relation priming, as shown in Figure 3-B and 3-C. During the N200 period, although not reach a significant level, the relation was activated to a certain extent at the three levels of NU, MS, and HS, which was manifested in the decrease in the amplitude of RS relative to RD. Seen from the extent of decline of amplitude, MS and NU have a certain degree of activation in the early stage of 200ms. The activation has increased over time and then has reached a very significant level in the N400 window. But at the HS level, there was almost no decrease in the N200 period; that is, there was no activation of relation, but in the N400 period, this activation also reached a very significant level, indicating that the relation associated with head activated later relative to MS and NU. In the N200 time window, the relation not worked at the HS level (that is, RS’s amplitude not declined relative to RD). However, its overall activation level is relatively high (amplitude is larger relative to MS and NU), indicating that HS has a relatively high activation level in an earlier period.

Moreover, HS has the shortest response time (MS: 794; HS: 771; NU: 850), indicating that its recognition is the fastest. It shows that although the head is the last morpheme, it plays a more important role in whole word processing than the first morpheme. We speculate that the head represents the conceptual category to which the entire word belongs. After it is activated, the approximate meaning of the whole word has been determined. Therefore, in the N200 period, the activation of the head inhibits the activation of relation associated with it until the later stage of word processing (N400 period) when more precise and detailed information needs to be extracted the relation associated with the head is activated. Ji’s results also support our explanation (Ji & Gagné, 2007). In his experiment, modifier or head appeared 350ms before the other constituent to determine the degree to which the relation is related to the two constituents. When the modifier appears 350ms before the head, the relation associated with the head is still active. But, when the head appears 350ms before the modifier, the relation associated with the modifier does not work, indicating that the head plays a more critical role than the modifier in processing Chinese compounds. When the head is pre-activated, the meaning of the entire word has been determined, and the relation associated with the modifier is no longer critical.

Then we combined the time course revealed by the ERP to explain the behavior results. It can be seen from Figure 3-A that, for the three morpheme levels, the amplitude reduction in RS relative to RD is the largest at the MS level, followed by the NU, and almost zero at the HS. It indicated that in the modifier-repeated conditions, relation plays the largest role in whole word processing. So the relation priming in the modifier-repeated condition was the first to be discovered by researchers (Gagné, 2000, 2001; Gagné & Shoben, 1997, 2002). In combination with the HS level results, we speculate that, when there is morpheme repetition, regardless of whether it is modifier or head, the relation is easier to extract. Head-repeated condition is an exception, in which relation priming is inhibited by constituent priming, so no effect was manifested. In the no morphemes repeated condition, the relation is more difficult to extract because subjects need to extract the relation of prime without the help of specific concepts and then apply it to target recognition. This process is relatively complicated, so the relation priming shown is smaller than that of MS, and the reaction time is longer.

Therefore, the relation activation pattern of the three levels of morphemes is relation activated at a very early stage (about 200ms), and morphological processing also occurs in this period (marked by the central-parietal N200). It indicates that the activation of the relation is almost simultaneous with the recognition of the constituent morphemes. In the earlier stage, the degree of activation is relatively small. As the word processing progresses, the activation gradually increases. Until the N400 period, the semantic composition begins after the semantic extraction of the constituent morphemes completed. There are only exceptions in HS. It is equivalent to a one-step success. The activation of constituents (i.e., head) is enough to activate the approximate meaning of the whole word. The relation only started to work when the details need to be modified later.

In our previous experiment, only the modifier-repeated condition was used (Jia et al., 2013). The relation activation during the N200 time window (although it was not significant) was ignored, leading to the conclusion that relation activation occurs in the semantic level. In the present experiment, activations in N200 happened at all three levels of morphemes. Although not reached significant levels, we couldn’t ignore this trend. It shows that relations started to activate at the lexical level. Since the morphology of the two constituents is also processed in this stage (Zhang et al., 2012), we have reason to speculate that the two constituent morphemes and relation, as semantic composition elements, are processed almost simultaneously. This is supported by the current literature (Gagné & Shoben, 2002; Raffray, Pickering, & Branigan, 2007).

These studies regard relations as a node on the semantic network. Like the constituent morphemes, they can be used independently as elements of composition. Morphemes can be associated with thousands of relations, and relations can also be related to thousands of morphemes. Whether primed by morphemes or relations, one is pre-selected from thousands of candidates, so the earlier this process occurs, the more economical it is for cognitive processing. Our present results support this speculation, and this process started as early as about 200ms. Moreover, the earlier the relation is activated, it will serve as a kind of micro-context, helping to determine the constituent morphemes’ meanings, especially in the case of ambiguous compounds.

Whether muti-morpheme words are processed as a whole or decomposed into smaller units has always been the focus of debate (Badecker, 2007; Taft & Forster, 1975). The discovery of the independent representation of relation in the present study adds new evidence for the view of decomposition. However, the materials used in the current experiment are transparent words with relatively low frequency. For compounds with unusually high frequency and compounds that cannot be decomposed literally, whether it is decomposed and processed in this way is worthy of further study in the future.

## 5. Conclusion

In this study, using Chinese two-morpheme compounds as material, we adopted a priming paradigm to investigate the relation priming of three morphemes level (modifier-repeated, head-repeated, and no repeated morpheme). In addition to the morpheme-repetition condition, we found that the relation priming can also occur without repeated morphemes, indicating that relation can be represented independently and does not have to be bound to a specific morpheme. The results on ERP shows that relation activated as early as about 200ms (N200). With the word processing progress, the activation continues to increase (N400), indicating that relation information plays an increasingly important role in the compound word’s recognition process. First, as contextual information, it helps to determine the meaning of the constituent morphemes. Then after the meaning access of the constituent morphemes, they begin to play a role in the semantic composition process. As the first ERP study to discover the independent representation of the relation information, it also provided a new perspective for compound words’ semantic access process by depicting the time course of relation information.

## Notes

**The work described in this paper was supported by the Key project of national key R & D project(No. 2017YFC1703303).**

### Competing Interest Statement

The authors have declared no competing interest.

